# Do Regional Brain Volumes and Major Depressive Disorder Share Genetic Architecture?: a study of Generation Scotland (n=19,762), UK Biobank (n=24,048) and the English Longitudinal Study of Ageing (n=5,766)

**DOI:** 10.1101/059352

**Authors:** Eleanor M. Wigmore, Toni-Kim Clarke, Mark J. Adams, Ana M. Fernandez-Pujals, Jude Gibson, David M. Howard, Gail Davies, Lynsey S. Hall, Yanni Zeng, Pippa A. Thomson, Caroline Hayward, Blair H. Smith, Lynne J. Hocking, Sandosh Padmanabhan, Ian J. Deary, David J. Porteous, Kristin K. Nicodemus, Andrew M. McIntosh

## Abstract

Major depressive disorder (MDD) is a heritable and highly debilitating condition. It is commonly associated with subcortical volumetric abnormalities, the most replicated of these being reduced hippocampal volume. Using the most recent published data from ENIGMA consortium’s genome-wide association study (GWAS) of regional brain volume, we sought to test whether there is shared genetic architecture between 8 subcortical brain volumes and MDD. Using LD score regression utilising summary statistics from ENIGMA and the Psychiatric Genomics Consortium, we demonstrated that hippocampal volume was positively genetically correlated with MDD (r_G_=0.46, *P*=0.02), although this did not survive multiple comparison testing. None of other six brain regions studied were genetically correlated and amygdala volume heritability was too low for analysis. We also generated polygenic risk scores (PRS) to assess potential pleiotropy between regional brain volumes and MDD in three cohorts (Generation Scotland; Scottish Family Health Study (n=19,762), UK Biobank (n=24,048) and the English Longitudinal Study of Ageing (n=5,766). We used logistic regression to examine volumetric PRS and MDD and performed a meta-analysis across the three cohorts. No regional volumetric PRS demonstrated significant association with MDD or recurrent MDD. In this study we provide some evidence that hippocampal volume and MDD may share genetic architecture, albeit this did not survive multiple testing correction and was in the opposite direction to most reported phenotypic correlations. We therefore found no evidence to support a shared genetic architecture for MDD and regional subcortical volumes.

## Introduction

Major depressive disorder (MDD) is a debilitating condition that accounts for a large proportion of disease burden world-wide^1^. It is a complex disorder that is influenced by both genetic and environmental factors with a heritability of approximately 37% estimated from twin studies^2^. Two recent genome-wide association studies (GWAS) identified two loci in MDD^3^ and 15 loci in self-reported depression^4^ of genome-wide significance. Nevertheless, the majority of MDD’s heritability is unaccounted for by currently identified variants and the mechanisms leading from gene to clinical phenotype remain elusive.

Reports of lower brain volumes in cross-sectional studies are common in MDD, but small sample sizes have led to poorly replicated results. Enhancing Neuro-Imaging Genetics through Meta-Analysis (ENIGMA) completed a large MDD case-control meta-analysis of subcortical volumes (n=8,927) demonstrating a significant association between MDD and reduced hippocampal volume (Cohen’s *d*=−0.20)^5^. Numerous other studies have also demonstrated a link between hippocampal reduction and MDD and it is one of the most robustly associated regions^6^. Other brain regions have shown limited and sometimes contradictory evidence for association with MDD. Smaller amygdala volume has been associated with depressive symptoms^7, 8^ and MDD status^9^, however larger amygdala volume has also been associated with the disorder^10^. A 2013 meta-analysis concluded that, as well as hippocampus, smaller putamen and thalamus volumes were associated with late life MDD, although fewer studies have examined these regions^11^. Additionally, smaller caudate nucleus volumes have also been associated with MDD in a meta-analysis^12^. The nucleus accumbens has not been widely associated with MDD status but a smaller volume has been implicated in the lethality of suicidal acts within mood disorder sufferers^13^. Pallidum volume and intracranial volume (ICV) have not been associated with MDD in any meta-analysis to date.

Subcortical structural volumes are known to be influenced by both genetic and environmental factors and have been demonstrated to be moderately to highly heritable ranging from 0.44 to 0.88^14^. The previously reported lower brain volumes in MDD and the relatively high heritability of these structures means they could be of interest as an intermediate phenotype^14^ potentially identifying the risk conferring loci as well as the underlying mechanisms of MDD. A GWAS of regional brain volumes has recently been completed by the ENIGMA Consortium^15^, providing an important opportunity to examine the genetic overlap between subcortical brain volumes and MDD. Overlap between genes involved in MDD and subcortical regions have been explored previously. The majority of studies have focused on candidate genes, such as the serotonin transporter (5-HTTLPR), and findings have often been contradictory^16^.

In this current study, we sought to examine whether the genetic architecture of MDD is shared with that of multiple subcortical brain regions. We employed two techniques; the first, LD score regression^17, 18^, estimates the genetic correlation between these traits using summary statistics from the ENIGMA and PGC consortia. The second method, polygenic risk scoring^19^, utilises ENIGMA summary statistics to generate individual level polygenic profile scores of each brain region’s volume. We then calculated the association of polygenic scores with MDD status in three cohorts, adjusting for confounds on an individual subject level, and combined them in a meta-analysis.

## Methods and Materials

### Cohort Descriptions and genotyping

#### Generation Scotland: Scottish Family Health Study (GS:SFHS)

GS:SFHS is a family based cohort with phenotypic data for 24,080 participants (mean age=47.6, s.d.=15.4) of which 20,032 participants had genotype data. Recruitment for this cohort has been described previously^20^. Diagnosis of MDD was made using the structured clinical interview for DSM-IV disorders (SCID) if individuals screened positive during interview questions (n=19,762, cases=2,643)^21^. Bipolar disorder patients (n=76) were excluded from this study. Information on MDD episodes and age of onset was also included in the SCID and therefore recurrent MDD and duration of MDD could be inferred (further details are given in the supplementary materials).

Details of the DNA extraction for GS:SFHS have been previously described^22^. Genotyping was completed at the Wellcome Trust Clinical Research Facility Genetics Core, Edinburgh (www.wtcrf.ed.ac.uk) using the Illumina HumanOmniExpressExome -8v1.0 Beadchip and Infinium chemistry^23^ and processed using GenomeStudio Analysis Software v2011.1. Quality Control (QC) utilised the inclusion threshold as follows; missingness per individual <1%, missingness per single nucleotide polymorphism (SNP) <1%, Hardy-Weinberg Equilibrium (HWE) p-value >1x10^-6^, minor allele frequency (MAF) >1%. 556,705 SNPs and 19,994 individuals passed QC criteria.

#### UK Biobank

UK Biobank is an open resource cohort with phenotypic data for 502,664 (mean age=56.5, s.d.= 8.1) between the ages of 40-69 recruited within the United Kingdom between 2006-2010 and genotype data available for 152,734 participants. Our study was conducted under UK Biobank application 4844. Study design and recruitment has been described previously^24^ but, in brief, participants were asked to complete a touchscreen questionnaire and additional data was collected by nurse interview. MDD status was based upon putative MDD phenotype defined by Smith et al., (2013)^25^ (n=24,048). Participants with mild depressive symptoms were removed based on this definition and self-reported bipolar disorder participants (n=1,211) were excluded. Information on MDD episodes and age of onset was also available therefore recurrent MDD and MDD duration was inferred (further details are given in the supplementary materials). Subcortical volumes for nucleus accumbens, amygdala, caudate nucleus, hippocampus, pallidum, putamen and thalamus were measured by T1-weighted structural imaging. The UK Biobank imaging protocol has been described elsewhere (http://biobank.ctsu.ox.ac.uk/crystal/refer.cgi?id=1977). The mean of the sum of left and right volume was taken for each subcortical region. ICV was generated by the sum of white matter, grey matter and ventricular cerebrospinal fluid volumes. Imaging data for the eight structures was available for 4,446 participants of which 968 had genetic data available.

Genotyping was completed utilising two Affymetrix arrays; BiLEVE (n=49,979) and the UK Biobank Axiom (n=102,750). Details have been described previously^26^. Initial genotyping QC was performed by UK Biobank^27^. Additional filtering was then applied to participants with poor heterozygosity or missingness, QC failure, non-British White ancestry, gender mismatch, genotype missing > 2%, and relatedness within UK Biobank and to the GS:SFHS sample (r > 0.0442, n=35,752) and ELSA sample (in the meta-analysis with all 3 cohorts). SNPs inclusion criteria were HWE p > 1x10^-6^ and MAF > 1%. 731,536 SNPs and 152,735 individuals passed QC criteria.

#### English Longitudinal Study of Ageing (ELSA)

ELSA is a prospective cohort study of health and ageing collected in 2002 with six follow-up waves taken at two-year intervals. At wave 1 (baseline) phenotypic data were available for 12,003 (mean age=63.9, s.d.=10.7) and genotypic data available for 7,452 participants. Details of this cohort have been described previously^28^. MDD status in this study was defined using a shortened form of the Centre of Epidemiological Studies – Depression scale (CES-D scale) (completed by 5,752 participants with genomic data). This consisted of 8 questions, rather than the original 20, with a “no”/”yes” response which was converted to a binary 0/1, respectively, although positive questions i.e. “During the past week, were you happy?”, were scored in reverse; 0 being “yes” and 1 being “no”. After summing the scores, a dummy variable of MDD status was classified as those with a score of 4 or above, as in previous studies^29^. Self-reported “manic depressive” (n=41) individuals were excluded.

Genotyping was completed in 2013/14 on 7,452 participants on the Illumina Omni 2.5-8 chip and QC was completed at the University College London Genetics Institute including exclusion of related individuals (r >= 0.2, n=109). Further QC was implemented using the same inclusion thresholds as used for GS:SFHS; SNP inclusion criteria were HWE p > 1x10^-6^ and MAF > 1%. >1.3 million SNPs and 7,230 individuals passed QC criteria.

### Linkage disequilibrium (LD) score regression

Genetic correlation of subcortical structures and MDD were measured using the LD score regression technique^17, 18^. In brief, this technique utilises GWAS summary statistics to estimate the SNP-based heritability of a trait and genetic correlation between traits, in this study we used summary data from ENIGMA and PGC. SNPs inclusion criteria were INFO > 0.9 and MAF > %1 (further details in supplementary materials).

Summary statistics for the regional brain volume GWAS completed by ENIGMA were downloaded from http://enigma.ini.usc.edu/enigma-vis/. The GWAS was completed on 11,840 participants on eight MRI volumetric measures; nucleus accumbens, amygdala, caudate nucleus, hippocampus, pallidum, putamen, thalamus and ICV^15^.

Summary statistics for the MDD GWAS completed by the MDD Working Group of the Psychiatric Genomics Consortium were downloaded from http://www.med.unc.edu/pgc/downloads. The study examined 9,238 MDD cases and 8,039 controls^30^.

### Polygenic Risk Scoring (PRS)

Construction of PRS was completed in PLINK software^31^, which has been previously described ^19^. Summary statistics were taken from the ENIGMA GWAS^15^ (details above) to construct weighted PRS using five *P* value thresholds: 0.01, 0.05, 0.1, 0.5 and 1, after SNPs underwent clumped-based pruning (r^2^=0.25, 300kb window). All five thresholds are reported in models of subcortical volume PRS predicting their respective volume in UK Biobank and the best predictive threshold was carried forward into models associating MDD status with each subcortical volume in all three cohorts. The *P* value thresholds carried forward were; nucleus accumbens: 0.01, amygdala: 0.1, caudate nucleus: 0.5, hippocampus: 0.01, ICV: 0.5, pallidum: 0.5, putamen: 0.1 and thalamus: 0.05. Scores for GS:SFHS, UK Biobank and ELSA were computed on the raw genotypes.

### Statistical Analysis

Mixed linear model analyses were completed in ASReml-R (http://www.vsni.co.uk/software/asreml/) for GS:SFHS with MDD status as the dependent variable and volume PRS fitted as the independent variable. The model was adjusted for age and sex with the first four principal components (PCs) fitted to control for population stratification. An additive matrix (expected relatedness derived from pedigree information) was fitted as a random effect to account for the family structure in GS:SFHS. Wald’s conditional F-test was used to calculate *P* values for all fixed effects and the variance explained was calculated by division of the difference in the sum of residual variance and additive genetic effect in the null model (without PRS) with the full model (with PRS). To adjust for the use of linear mixed regression models being applied to a binary dependent variable in a structured dataset, the fixed effects and standard errors from the linear model were transformed utilising a Taylor series approximation from the linear scale to the liability scale utilising a previously described technique^32^. Since hippocampal volumetric differences have been more closely associated with recurrent MDD and early illness onset^5, 33^, hippocampus PRS regression analyses were also run with recurrent MDD, number of episodes, MDD duration and age of onset as dependent variables (for further details see Supplementary Materials).

Logistic regression utilising generalised linear models in R version 3.2.3 (www.r-project.org) was used for both the UK Biobank and ELSA cohorts to test association between MDD and PRS of subcortical volumes. Models were adjusted for age, sex and the first 15 PCs (in UK Biobank) and first 4 PCs (in ELSA) to control for population stratification. Linear regression was used for models predicting subcortical structures and these were also adjusted for ICV. Hippocampus volume PRS was also examined with recurrent MDD, number of episodes, MDD duration and age of diagnosis for UK Biobank however this data was not available for ELSA (see Supplementary Materials).

In order to increase power, fixed effect meta-analysis, weighted by standard error of the beta values relating PRS scores to MDD was carried out using the ‘meta’ package (version 4.3-2)^34^ in R.

### Power analysis

Power analyses for the genetic correlations, calculated using LD score regression, were completed using the GCTA-GREML power calculator^35^. As LD score regression utilises summary statistics and GCTA the individual genotype data, true power is likely to be slightly lower however the GCTA-GREML power calculator gives a close estimate. Results of the power analysis are presented in **Supplemental Table S1**.

Simulations of 50% genetic correlation between the brain volume and MDD indicated we had power to detect an association in all regions. However, genetic correlation was found to be much lower for many of the regions and therefore we only had adequate power to detect a correlation between hippocampal volume and MDD (99%). For the remaining regions a power curve was conducted to demonstrate the size of the sample needed for sufficient power (**Figure 1** and **Supplemental Figure S1**). Results indicate that an additional ~15,000 sample increase in both ENIGMA and PGC samples would be needed to detect genetic correlation for Putamen and ICV at the genetic correlations reported in this analysis whilst nucleus accumbens would need a sample size increase of nearly 100,000 in both samples and pallidum significantly more than that.

**Figure 1.**
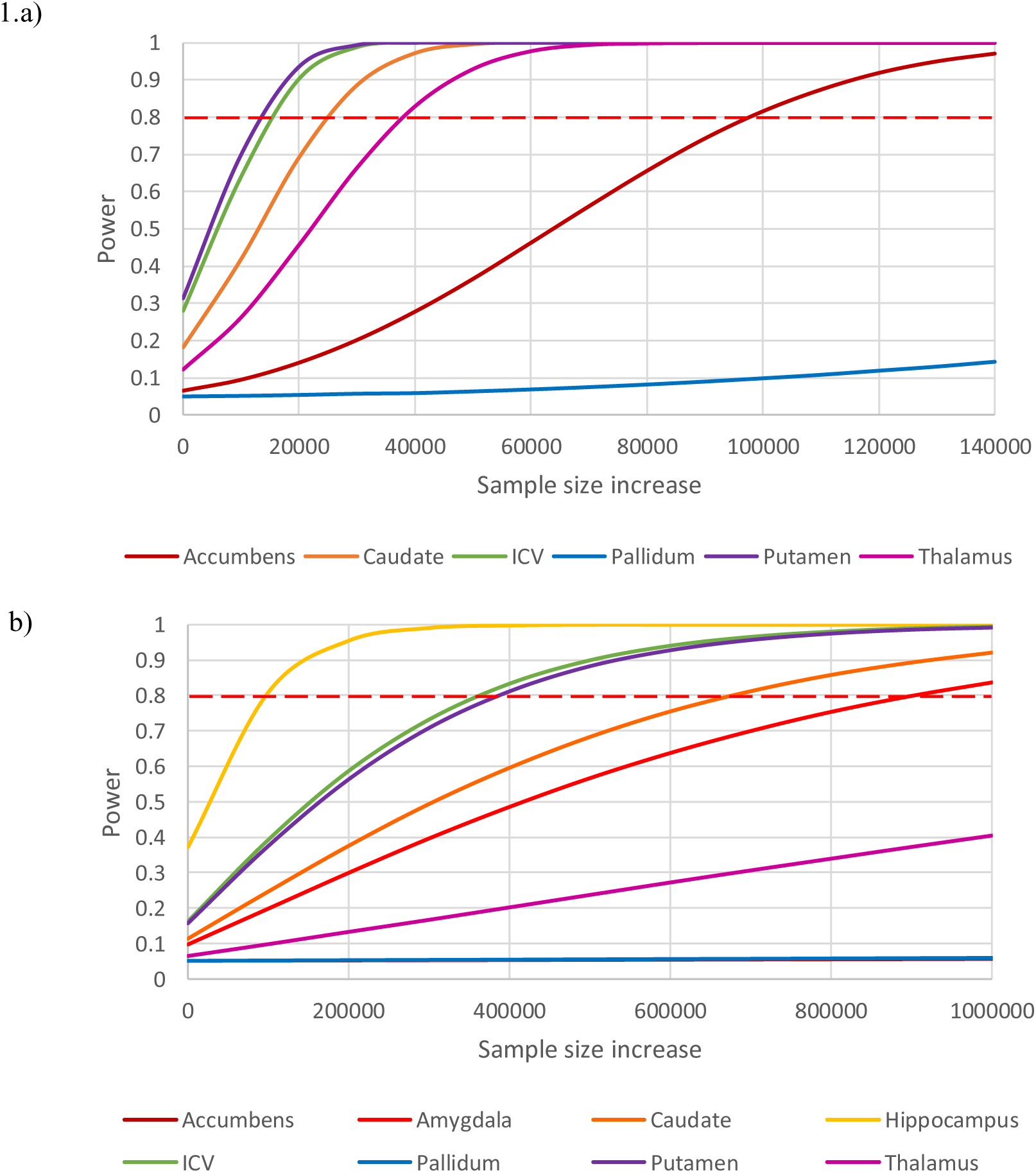
Power curves were calculated with starting point 0 as the sample size in our analysis. For the genetic correlation power analysis (a) sample sizes were increased equally in both samples and for MDD the proportion of cases and controls was kept constant. For PRS power analysis (b) the sample size for the training set (ENIGMA) was kept constant whilst the target set sample size was increased. Amygdala was assumed to have a covariance of 25% for the PRS power analysis. Hippocampus had adequate power in the genetic correlation analysis and therefore was not included in the power curve.

Power analysis of PRS were completed using AVENGEME^36^. Markers were assumed to be independent and 5% of SNPs were assumed to have an effect in the training sample. Genetic covariance values were taken from the LD score regression analysis, however, no value could be computed for amygdala therefore 3 hypothetical covariance’s were tested; 0.50, 0.25 and 0.10. Results of the power analysis are presented in **Supplemental Table S3**.

Despite the study number of 49,576 individuals, PRS was under-powered in all analyses. Highest power in the meta-analyses was hippocampus (37%). Low SNP heritability and low covariance between traits account for the low power in the meta-analyses. In the PRS analysis on their own trait, highest power was the putamen (23%) at a *P* value threshold of 1. In this analysis a small sample size of 968 individuals likely reduced power. We therefore conducted a power curve for both the meta-analysis and PRS in their own trait to indicate the sample size that would be necessary to have adequate power (**Figure 1** and **Supplemental Figure S2**). Power curves indicate that a sample increase of ~100,000 individuals in the target set would be sufficient power for hippocampus PRS associated with MDD, however nearly 900,000 for amygdala assuming a covariance of 0.25 and an increase of over 1 million for nucleus accumbens, pallidum and thalamus.

## Results

### Genetic correlation

Using LD score regression, we calculated SNP-based heritability estimates for the eight subcortical regions and MDD, utilising summary data from GWAS completed by ENIGMA^15^ and PGC^30^ respectively. The estimate of the SNP heritability for the amygdala was non-significant and therefore the amygdala was not included in any further analysis. SNP heritability estimates for the remaining subcortical volumes ranged from the SNP h^2^=0.0855 (s.e.=0.0438) for the nucleus accumbens to SNP h^2^=0.297 (s.e.=0.051) for the putamen (**Table 1**). MDD SNP heritability was calculated at 0.204 (s.e.=0.0386). Genetic correlation between each subcortical region and MDD was then calculated. Hippocampal volume demonstrated significant genetic correlation with MDD (r_G_=0.460, s.e.=0.200, *P*=0.0213) (**Table 1**), although this did not survive multiple testing correction (**Supplementary Table S3**). No other subcortical volume was genetically correlated with MDD.

**Table 1.**
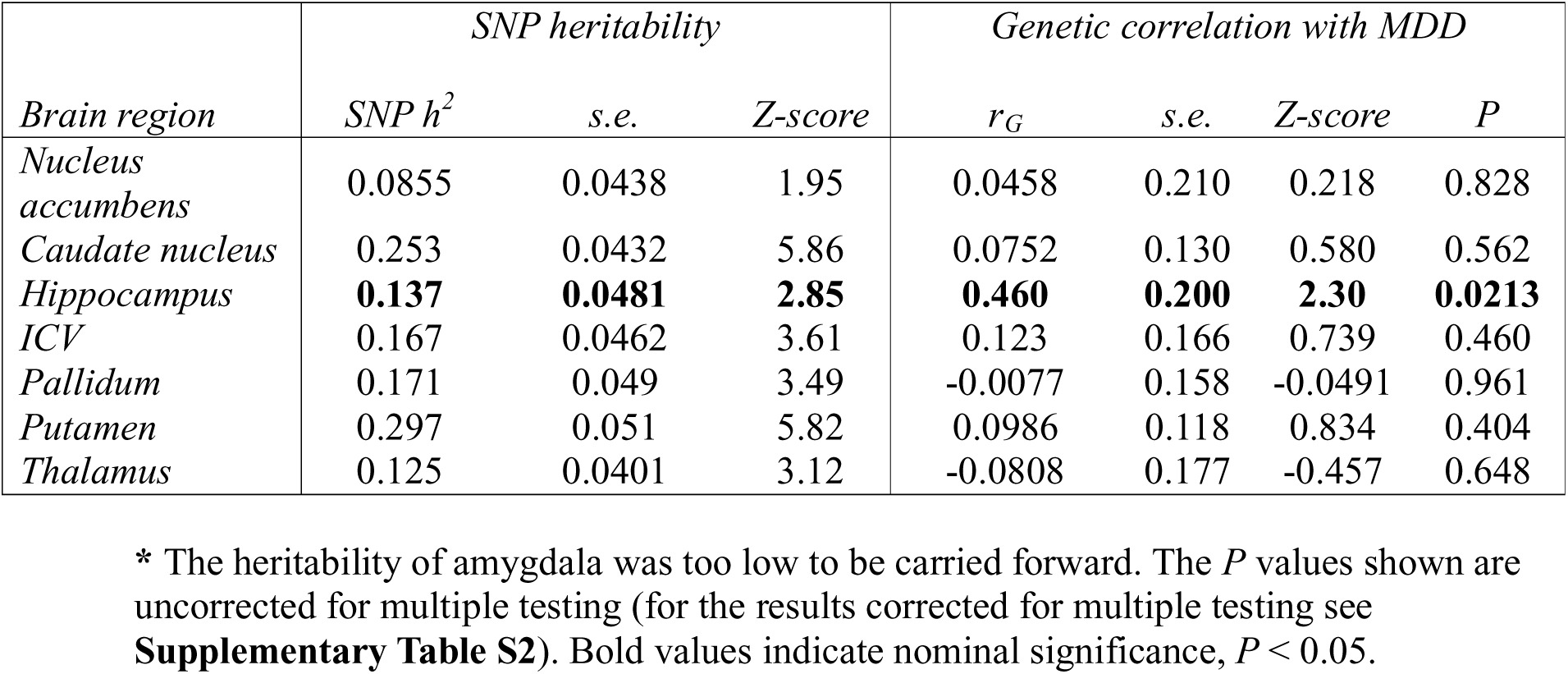
SNP-based heritability and genetic correlation of subcortical brain regions and MDD.

### Polygenic risk score (PRS)

Subcortical PRS were calculated in UK Biobank to examine the variance explained with respect to their own volume. PRS were positively associated with their respective volume in four of the eight structures across the 5 *P* value thresholds; caudate nucleus, ICV, putamen and thalamus. In addition, hippocampus was significantly associated at a *P* value threshold of 0.01 only. These results retained significance after multiple test correction across the 5 thresholds, however only raw *P* values have been reported. Nucleus accumbens, amygdala and pallidum PRS did not demonstrate any association with their respective volume. The variance explained by PRS was small for all volumes, with the largest reported in the caudate nucleus (R^2^=0.0102, beta=0.117, *P*=1.08x10^-4^) (**Figure 2** and **Supplementary Table S2**).

**Figure 2.**
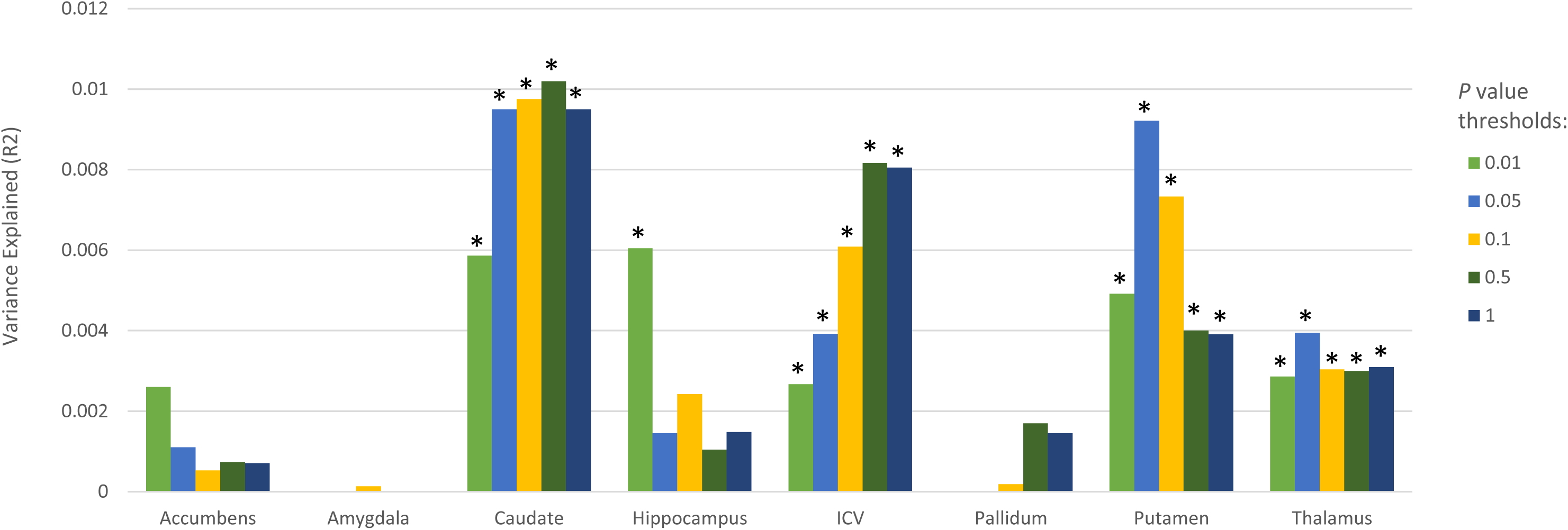
Significant *P* values (<0.05) are indicated with *. Nucleus accumbens, amygdala and pallidum PRS were not significantly associated with their respective volume at any threshold.

Structural PRS were selected at the threshold which best predicted its own volume (nucleus accumbens=0.01, amygdala=0.1, caudate nucleus=0.5, hippocampus=0.01, ICV=0.5, pallidum=0.5, putamen=0.1, thalamus=0.05) and tested for prediction of MDD status. No PRS for any volume was significantly associated with MDD status in any of the cohorts (**Table 2**). In order to increase power, we completed a meta-analysis of the summary association statistics from three cohorts. No evidence of heterogeneity was identified in any of the meta-analyses. We found no association between any structural PRS and MDD (**Figure 3a** and **Supplementary Fig. S3**). Association between hippocampal volume and recurrent MDD and early illness onset has been previously reported^5, 33^. We therefore examined MDD phenotypes in association with hippocampal volume PRS in GS:SFHS and UK Biobank, this data was not available for the ELSA cohort. There was no association between hippocampal PRS and recurrent MDD (OR=0.98, *P*=0.0850) (**Figure 3b**). Further, hippocampal volume PRS was not significantly associated with number of episodes (beta=-0.00390, *P*=0.425), MDD duration (beta=-0.00110, *P*=0.414) or age of onset (beta=0.0142, *P*=0.291) (**Supplementary Fig. S4**).

**Table 2.** Mixed model analysis of Subcortical volumetric PRS and MDD in all 3 cohorts.

| *PRS* | **Study** | **P value** | **beta** | **s.e.** |
| --- | --- | --- | --- | --- |
| *Nucleus accumbens* | **GS** | 0.485 | -0.0181 | 0.0218 |
|  | **UKB** | 0.284 | -0.0151 | 0.0141 |
|  | **ELSA** | 0.659 | 0.0175 | 0.0395 |
| *Amygdala* | **GS** | 0.327 | -0.0285 | 0.0217 |
|  | **UKB** | 0.962 | 0.000694 | 0.0146 |
|  | **ELSA** | 0.208 | -0.0500 | 0.0398 |
| *Caudate nucleus* | **GS** | 0.246 | 0.0197 | 0.0224 |
|  | **UKB** | 0.396 | -0.0135 | 0.0159 |
|  | **ELSA** | 0.114 | -0.0630 | 0.0398 |
| *Hippocampus* | **GS** | 0.782 | -0.00299 | 0.0211 |
|  | **UKB** | 0.601 | -0.00732 | 0.0140 |
|  | **ELSA** | 0.571 | -0.0222 | 0.0392 |
| *ICV* | **GS** | 0.395 | 0.0179 | 0.0225 |
|  | **UKB** | 0.280 | -0.0160 | 0.0148 |
|  | **ELSA** | 0.0995 | 0.0652 | 0.0396 |
| *Pallidum* | **GS** | 0.752 | -0.00808 | 0.0221 |
|  | **UKB** | 0.250 | 0.0178 | 0.0155 |
|  | **ELSA** | 0.658 | 0.0176 | 0.0398 |
| *Putamen* | **GS** | 0.303 | -0.0246 | 0.0218 |
|  | **UKB** | 0.215 | 0.0231 | 0.0186 |
|  | **ELSA** | 0.695 | -0.0161 | 0.0410 |
| *Thalamus* | **GS** | 0.451 | -0.0192 | 0.0219 |
|  | **UKB** | 0.836 | 0.00293 | 0.0142 |
|  | **ELSA** | 0.222 | -0.0482 | 0.0395 |

\* Best *P* value threshold for PRS were carried forward in this analysis; Nucleus accumbens=0.01, Amygdala= 0.1, Caudate nucleus=0.5, Hippocampus=0.01, ICV=0.5, Pallidum=0.5, Putamen=0.1, Thalamus=0.05. No PRS demonstrated a significant association with MDD in any cohort.

**Figure 3.**
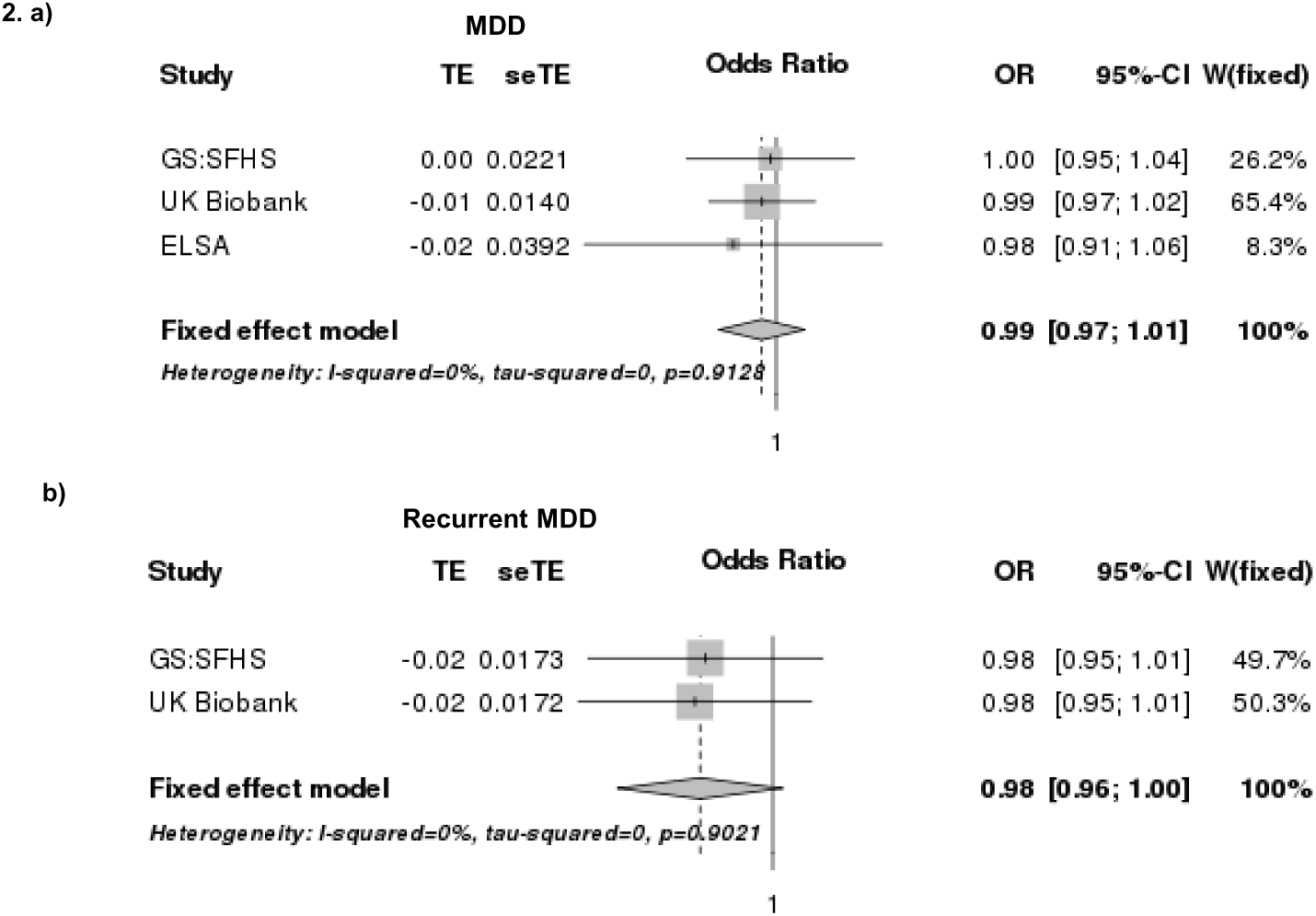
Both plots demonstrate a negative correlation with MDD and recurrent MDD with no heterogeneity between cohorts but neither plot reaches statistical significance. TE; treatment effect (regression beta’s); seTE, standard errors; OR, odds ratio; CI, confidence intervals; W(fixed), weight of individual studies in fixed effect meta-analysis.

## Discussion

Previous studies have reported phenotypic associations between brain volumes and MDD. In this study we investigated whether there was evidence of shared genetic architecture between subcortical brain volumes and MDD. Genetic correlations between regional brain volume and MDD show that hippocampal volume and MDD are partially influenced by common genetic variants (r_G_=0.46, s.e.=0.200, *P*=0.0213), although this did not survive correction for multiple testing. No other brain regions’ volume showed evidence of shared genetically aetiology with MDD. This sample size was adequate to detect an association at 50% genetic correlation, however, at the genetic correlation reported in this study we were underpowered in all other brain volumes (excluding hippocampus). Further analysis is needed utilising larger sample sizes (a minimum of 15,000 increase in both samples) to draw confirmatory conclusions in other regions. A meta-analysis of data from three studies, totalling 49,576 individuals including 11,552 cases, found no evidence of association between any regional brain volume PRS and MDD, including the hippocampus. Since previous neuroimaging evidence suggests that decreased hippocampal volumes could occur as a consequence of recurrent depressive episodes and early illness onset^5, 33^, we examined hippocampal volume PRS in association with recurrent MDD, number of episodes, MDD duration, and age of onset but we observed no significant associations.

The genetic correlation reported between total hippocampal volume and MDD is novel, so far as any of the authors are aware, however a genetic correlation has been reported between recurrent MDD and right hippocampal volume. Mathias et al., (2016) demonstrated significant negative correlation measured via linkage analysis in a sample of 893 individuals^37^. Our finding, however, was a positive correlation which was not significant after correcting for multiple testing. Despite the PRS meta-analysis being the largest analysis to date examining genetic scores for brain volume and MDD it was severely underpowered therefore we can draw no confirmatory conclusions about the genetic overlap between any structure and MDD from this analysis. The apparent discrepancy between PRS and our finding LD score regression is possibly due to this lack of power, however, several other reasons present possible explanations. Firstly, linkage disequilibrium (LD) information is utilised in LD score regression to measure genetic correlation, however in the process of PRS calculation, all SNPs are pruned to a much smaller independent set. Previous simulation studies have demonstrated that predictive capabilities of PRS are greatly enhanced when utilising LD information^38^. This implies that LD pruning may be removing causal SNPs and those more closely tagging causal variants, resulting in a loss of information and predictive accuracy. Secondly, each dataset had a different MDD definition; GS:SFHS utilised the SCID, ELSA MDD was defined utilising the CES-D and UK Biobank MDD was generated using self-reported information. Whilst the PGC MDD definition most closely matches that of GS:SFHS MDD, the GS:SFHS sample was population based rather than identified from a clinically ascertained samples. The observed lack of replication using the described methodologies may therefore be due to factors related to ascertainment differences. Thirdly, PRS have not explained a large proportion of variance within their own trait in almost any analysis conducted to date, with the possible exception of schizophrenia. The PRS that explained most variance in hippocampal volume only explained 0.621%, however, this analysis was also hindered by lack of power.

We conclude that hippocampal volume and MDD may share common genetic factors, although this result did not withstand multiple test correction. Animal models have previously demonstrated that increased stress can drive decreased hippocampal neurogenesis (and therefore increased atrophy)^39^ and this reduced neurogenesis can lead to depressive-like symptoms^40^. Stress is a well-established environmental risk factor associated with MDD^41^ and the inhibition of glucocorticoid receptors has been shown to normalise hippocampal neurogenesis^42^ and relieve symptoms in psychotic major depression^43^. Furthermore, increased duration of depression has also been related to more pronounced hippocampal reductions^44^. Our results however indicated a positive genetic correlation suggesting that genetic variants determining *larger* hippocampal volume may be risk factors for MDD. If this is the case, it is possible that an intermediary environmental process may be the reason for the deviation from previous literature. Hippocampal volume has been demonstrated to be more highly impacted by the environment than other brain regions^45^ and decreased hippocampal volume is associated with many environmental factors e.g. stress^41^, increased exercise^46^ and jet lag^47^. It is therefore possible that the previously reported decreased hippocampal volume associated with MDD is a due to multiple episodes of depression and that this positive genetic correlation is due to a role in MDD susceptibility earlier in brain development. Given the previous literature linking both hippocampal structure and MDD to stress, it is possible that gene-environment interactions (GxE) could further explain their correlation.

Hippocampal volume changes are also widely associated with other psychiatric disorders such as schizophrenia. A similar analysis that examined the genetic correlation between subcortical volumes and schizophrenia found no significant correlations^48^. This is suggestive that the genetic correlation observed could be specific to hippocampal volume in MDD. However, these results are only indicative of a genetic correlation between the two traits and further research would be necessary to provide confirmative evidence and the directionality of any causal relationships.

Subcortical volume PRS were not associated with their own volume in three out of the eight structures and was only associated with hippocampal volume at one threshold. This is likely due to the analysis being underpowered to detect an association in a sample size of 968 participants. Power of the PRS is also limited by the size of the initial ENIGMA GWAS (n=11,840), larger discovery sample sizes greatly improve the accuracy of PRS^49, 50^. Of the PRS that were associated with their phenotype, the largest proportion of variance explained was 1% with the majority predicting ~0.6%. The proportion of variance explained is therefore very low although this is fairly common in PRS studies^51^ with one of the largest explained variance by PRS reported in schizophrenia (~7% on the liability scale)^50^. It is therefore perhaps unsurprising that there was no significant association with MDD in the PRS which were not associated with their own phenotype.

This study has other notable limitations; for instance, this study only explored the effects of common genetic variants and it may be important to examine rarer variants to generate a more complete picture of their genetic architecture. PGC GWAS, despite being one of largest GWAS for MDD (~17,000), is potentially still too small to have adequate power to detect common variants^30^, likewise the ENIGMA study too may have insufficient power. The lower heritability, higher prevalence and likely heterogeneity of MDD results in less precise estimates of marker weights from GWAS^52^, decreasing the power to detect genetic correlations with other phenotypes. Larger genome-wide analysis would be necessary to generate confirmatory conclusions. The estimates for SNP heritability, calculated using LD score regression, were lower than have been previously described^53^. LD score regression has been utilised previously to calculate SNP heritability of subcortical volumes using the ENIGMA summary data with similar low estimates reported^48^.

Despite these limitations, we provide some evidence of a genetic correlation between hippocampal volume and MDD, however, we could not demonstrate an association utilising PRS techniques. Limited power, low explanation of variance and loss of LD information were notable limitations in our PRS analysis. Whilst we conclude; that the relationship between hippocampal volume and MDD could be in part driven by genetic factors, we think the most important outcome for the current study is in the planning for future studies. Sample sizes of ~150,000 individuals will be needed to have sufficient statistical power (>0.8) to detect shared genetic architecture between MDD and hippocampal volume using PRS, using datasets similar to the one studied. The other regional brain volumes ranged from needing an additional sample size of ~400,000 to in excess of 1 million individuals. Alternatively, further studies may utilise data from further releases of the ENIGMA consortium, including larger numbers of participants and more accurately determined SNP effect sizes.

## Acknowledgements

This investigation was supported by the Wellcome Trust 104036/Z/14/Z (STRADL, Stratifying Resilience and Depression Longitudinally). Generation Scotland received core funding from the Chief Scientist Office of the Scottish Government Health Directorate CZD/16/6 and the Scottish Funding Council HR03006. We thank all families, practitioners and the Scottish School of Primary Care involved in the recruitment process as well as the entirety of Generation Scotland team; interviewers, computer and laboratory technicians, clerical workers, research scientists, volunteers, managers, receptionists, healthcare assistants and nurses. We are grateful towards the Dr Mortimer and Theresa Sackler foundation for the financial support for this work. This research has been conducted using the UK Biobank resource and we would therefore like to thank all participants and coordinators in this cohort.'Samples from the English Longitudinal Study of Ageing DNA Repository (EDNAR), which receives support from the National Institute on Aging (NIA) and the Economic and Social Research Council (ESRC), were used in this study. We thank contributors and the ELSA participants. IJD is supported by MRC and BBSRC funding to the University of Edinburgh Centre for Cognitive Ageing and Cognitive Epidemiology (MR/K026992/1).

